# Immunohistochemical and ultrastructural analysis of the maturing larval zebrafish enteric nervous system reveals the formation of a neuropil pattern

**DOI:** 10.1101/555094

**Authors:** Phillip A. Baker, Matthew D. Meyer, Ashley Tsang, Rosa A. Uribe

**Affiliations:** Biosciences Department, MS 140, Rice University 6100 Main Street, Houston Texas 77005, USA; Shared Equipment Authority, MS 100, Rice University 6100 Main Street, Houston Texas 77005, USA

**Author notes:** email: Correspondence should be addressed to R.A.U.

## Abstract

The gastrointestinal tract is constructed with an intrinsic series of interconnected ganglia that span its entire length, called the enteric nervous system (ENS). The ENS exerts critical local reflex control over many essential gut functions; including peristalsis, water balance, hormone secretions and intestinal barrier homeostasis. ENS ganglia exist as a collection of neurons and glia that are arranged in a series of plexuses throughout the gut: the myenteric plexus and submucosal plexus. While it is known that enteric ganglia are derived from a stem cell population called the neural crest, mechanisms that dictate final neuropil plexus organization remain obscure. Recently, the vertebrate animal, zebrafish, has emerged as a useful model to understand ENS development, however knowledge of its developing myenteric plexus architecture was unknown. Here, we examine myenteric plexus of the maturing zebrafish larval fish histologically over time and find that it consists of a series of tight axon layers and long glial cell processes that wrap the circumference of the gut tube to completely encapsulate it, along all levels of the gut. By late larval stages, complexity of the myenteric plexus increases such that a layer of axons is juxtaposed to concentric layers of glial cells. Ultrastructurally, glial cells contain glial filaments and make intimate contacts with one another in long, thread-like projections. Conserved indicators of vesicular axon profiles are readily abundant throughout the larval plexus neuropil. Together, these data extend our understanding of myenteric plexus architecture in maturing zebrafish, thereby enabling functional studies of its formation in the future.

## Introduction

The enteric nervous system (ENS) consists of a multi-series web of thousands of interconnected ganglia that form nerve plexuses spanning circumferentially within the muscle walls of the entire gastrointestinal (GI) tract. Enteric ganglia are composed of enteric neuron subtypes and glial cells that communicate with each other to modulate neural reflexes of the gut^1^. As the largest component of the peripheral nervous system, the ENS enables the GI tract to perform critical life functions such as peristalsis, gut hormone secretions, and water balance during gut homeostasis^1^. The ENS is capable of autonomous reflex activity, separate from control of the central nervous system (CNS), and thus has been known as the “second brain”. Intense research has focused on ENS formation from its embryonic stem cell progenitor population, the neural crest (NCC), however knowledge of the cellular and molecular mechanisms that underpin its neural plexus manifestation and patterning is comparatively understudied.

ENS ganglia are derived from NCCs that originate from the dorsal neural tube during development^2–4^. In zebrafish, the ENS is derived from NCCs that originate from a post-otic area known as the vagal region^5–7^. Vagal NCCs migrate down the length of the primitive gut until they reach the distal-most hindgut, whereby they cease migration and terminally differentiate into neurons that are critical to forming neural circuits within the intestinal muscularis around the circumference of the entire gut tube by 3 days post fertilization (dpf)^8^. In all vertebrates, this neural circuit expansion leads to the development of a concentric neural plexus that continues to mature into the functional adult ENS^9^. In mammals, the ENS is comprised of two major layers: an outer myenteric plexus that is the first to form, followed by the inner submucosal plexus, both of which eventually display finer neuropil architecture over time^10,11^. Zebrafish possess a single myenteric plexus^12^. Currently, little is known regarding how the cellular constituents that make up the zebrafish myenteric plexus are patterned during development or whether a fine neuropil is evident along the gut length. Globally, understanding how myenteric plexus pattern is constructed during normal development will increase our ability to decipher how it is affected in pathological and diseased states, such as in Hirschsprung disease^13^ or Crohn’s Disease^14^. Therefore, a critical effort to study maturation of the enteric neuropil should be of great interest to developmental biologists, gastroenterology clinical experts and neuroscientists.

For decades, various studies have made great progress in increasing our understanding of the vertebrate ENS by investigating its formation and structure in various amniote systems^15^. However, microscopic study of these processes during early development has been hindered by the opaque nature of each organism and its amniotic development. Zebrafish have emerged as a powerful system to help build upon these studies and advance our understanding of ENS construction and function. Zebrafish larvae provide us with an incredible opportunity to do so during crucial developmental stages due to their transparent, and external embryonic and larval development^8^. In addition, zebrafish and mammalian gut organ systems are largely homologous—both exhibiting a foregut, midgut and hindgut—which allows us to glean important information and apply it to the understanding of more complex vertebrate development.

Previous research involving ultrastructural analysis of mammalian intestinal tissue has laid much of the foundation that supports our current understanding of neural and glial patterning within the ENS plexuses^16–18^. These ultrastructural investigations identified axon profiles and a large population of enteric glial cells that exist in the neuropil, outnumbering neurons within the adult ENS, that could be subclassified based on their morphology and location^16^. More recent studies involving immunohistochemical techniques in adult mice have supported these findings and found a 4-fold disparity in the number of glial to neurons. Additionally, they were able to make a distinction between various enteric glia based of the immunoreactivity of different glial markers^19^.

Previously, larval and adult zebrafish intestinal architecture has been examined using immunohistochemical and transmission electron microscopy (TEM) preparations^12^. While these studies have greatly improved our understanding of the zebrafish intestinal structure, particularly epithelial patterning, description of the myenteric plexus between the 6 dpf and adult stages, which may represent a key time when the ENS is patterned, is very meager. By utilizing a combination of high-resolution immunohistochemical and TEM techniques, here we report the presence of an enteric neuropil within the myenteric plexus of the maturing larval fish.

## Results

### I. Histology of the maturing larval zebrafish muscularis layer along the foregut, midgut and hindgut

To investigate organization of the zebrafish larval enteric plexus, we first sought to characterize general anatomy of the intestinal muscularis, within which the plexus rests, in 7 and 18 dpf fishes along the foregut, midgut and hindgut. At 7 dpf, the zebrafish gut is functionally developed and exhibits peristalsis activity^8,20^. By 18 dpf, the larval gut exhibits increased size and we postulated that it should exhibit a higher degree of maturation, when compared with the 7 dpf larval stage. Using transverse-sectioned histological analysis on plastic-embedded fishes and toluidine blue staining, we observe that the larval muscularis exists as a thin tissue layer in the outer circumference of the gut tube, lying directly outside of the intestinal epithelium (IE) at both time points (Fig.1, see areas in between dashed region). These observations agree with Wallace and Colleagues^12^ that have described the zebrafish muscularis layer as a relatively simple, thin layer along the gut tube. Within this tissue layer, along all levels of the gut, we observe that the muscularis contains a heterogeneous population of cells with toluidine blue stained nuclei at various densities, lightly stained cytoplasmic processes and varying relative total thicknesses throughout its circumference. Nuclei exhibit various shapes, ranging from circular and oval, to indented and very elongated, for example as in the hindgut (Fig.1C’,F’). While some nuclei are seen clustered together (Fig.1B,C), there are no apparent patterns. Overall, these histological observations indicate that the larval zebrafish muscularis does not undergo drastic changes in gross morphology between 7 and 18 dpf, however an increase in size of the muscularis is noted, along with growth of the fish.

**Figure 1:**
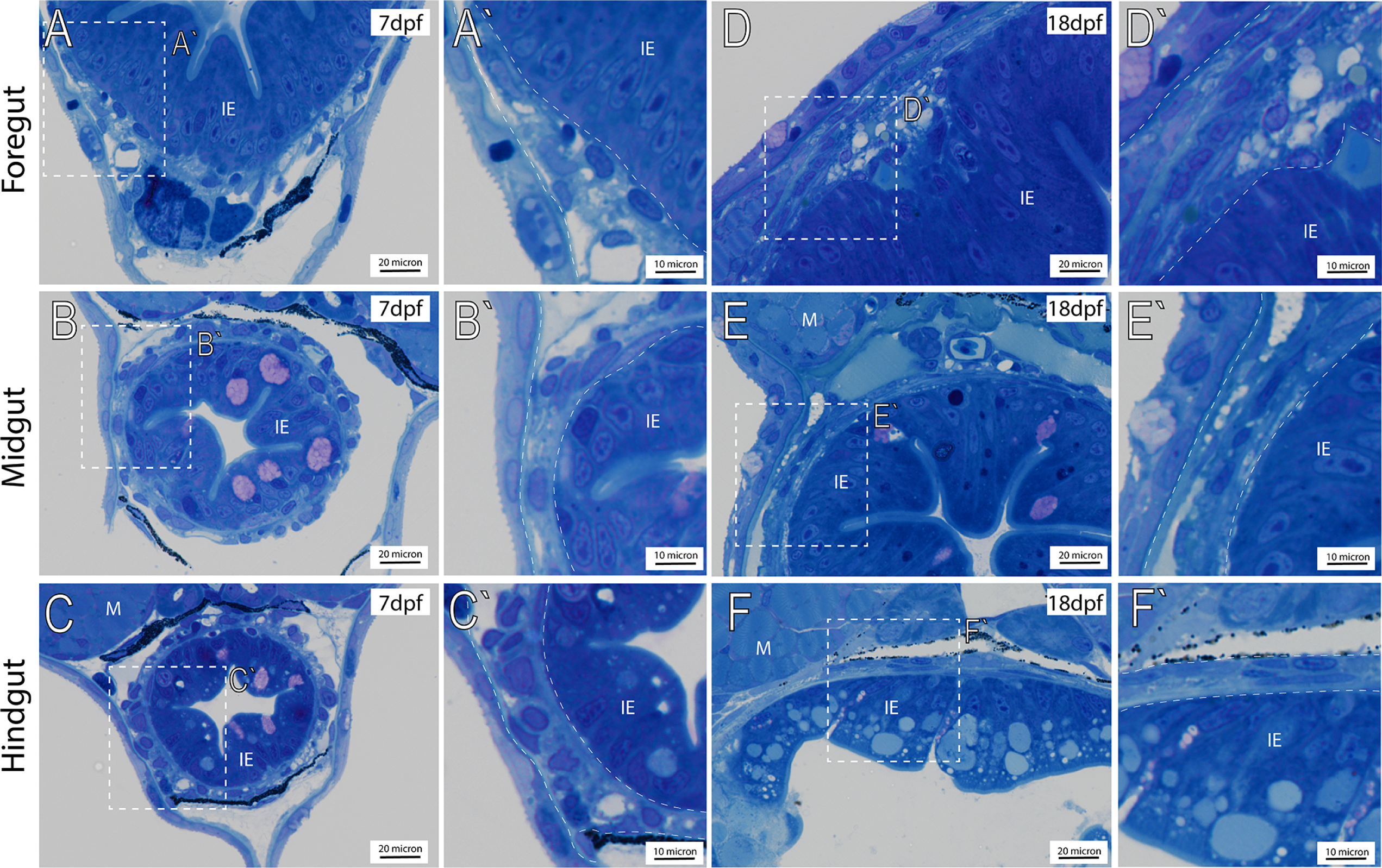
Transverse histological sections depict general anatomy of intestinal muscularis. Plastic embedded fish sectioned transversely and stained with toluidine blue at (A-C`) 7 dpf, and 18 dpf (D-F`), reveals the intestinal epithelium (IE) situated below trunk muscle (M). White-dashed box (A-F) corresponds to higher magnification panels (A`- F`) where intestinal muscularis (dashed lines).

### II. Axon and Glial cell localization in the maturing larval enteric plexus

To better understand spatial organization of the enteric plexus within maturing larvae, we examined transverse cryosections using immunohistochemistry to visualize the presence and circumferential distribution of axons and glia at 7 and 18 dpf, along the foregut, midgut and hindgut. To examine axonal projections, sections were processed using an antibody against Acetylated Tubulin (Acet-Tub), which has previously been used to mark neuron bodies and axons along the zebrafish gut tube^12,21–23^. To visualize glial cells within the zebrafish ENS at these stages, we utilized immunoreactivity of the zebrafish-specific Glial Fibrillary Acidic Protein (GFAP) antibody (Genetex GTX128741). GFAP is expressed in glial cells, including enteric glia^24^. This antibody labeled glial cells within the zebrafish CNS; in all sections there was a high degree of discrete immunoreactivity of GFAP throughout the brain (Fig.2A; arrows) and spinal cord (Fig.2B-D; arrows), revealing radial glia as previously observed^25^, confirming the ability of the antibody to label zebrafish glia.

**Figure 2:**
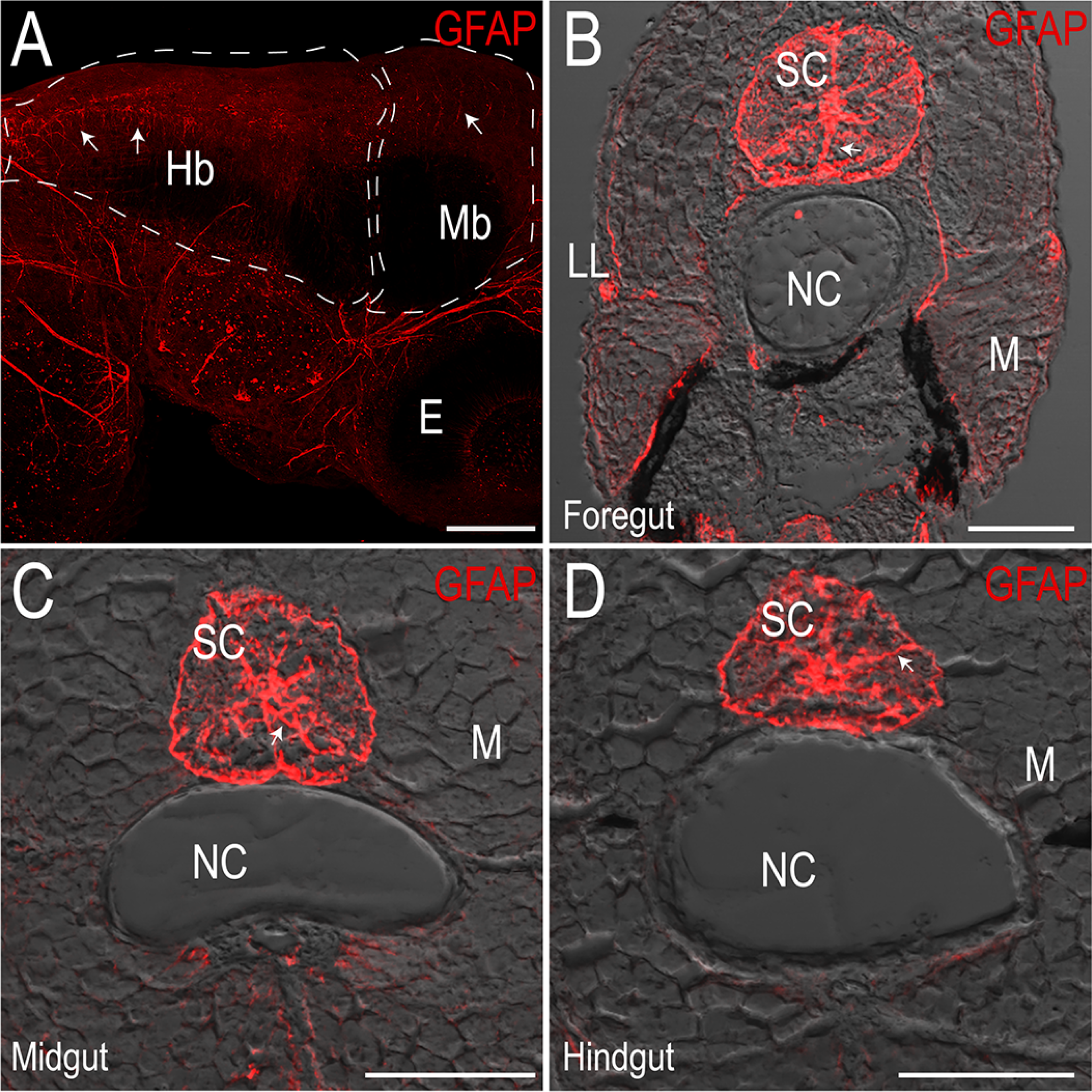
Zebrafish-specific Glial Fibrillary Acidic Protein (GFAP) antibody marks glia in the central nervous system. Maximum intensity projection confocal stack in (A) whole-mount immunofluorescence preparation of a 3 dpf larval fish showing GFAP^+^ cells in the hindbrain (Hb) and midbrain (Mb), scale bar: 100 microns, E: eye. (B-D) Maximum intensity projections of transverse cryosections of 18 dpf larvae at the level of the foregut (B), midgut (C) and hindgut (D) mark radial glia throughout the spinal cord (SC). LL: lateral line, NC: notochord, M: skeletal trunk muscle, scale bar: 50 microns.

In all sections of the gut, GFAP^+^ and Acet-Tub^+^ immunoreactive cellular processes span circumferentially around the outer layer of the gut tube, denoting the myenteric plexus of the ENS (Fig.3-5). Within the plexus, GFAP^+^ processes largely form an inner layer separating Acet-Tub^+^ axons from the IE, a general pattern observed across the foregut, midgut and hindgut, in both 7 dpf and 18 dpf larvae (Fig.3-5; Fig.9A,B). In particular, we noted that glial cells and axons immediately juxtapose one another along the gut circumference (Fig.3C,C’; arrowheads). Glial cell processes are long and thread-like, wrapping along the circumference of the gut tube to completely ensheathe it. Axons are also seen traversing between layers occasionally along all levels of the gut (Fig.3D’; Fig.4D’; Fig.5D).

**Figure 3:**
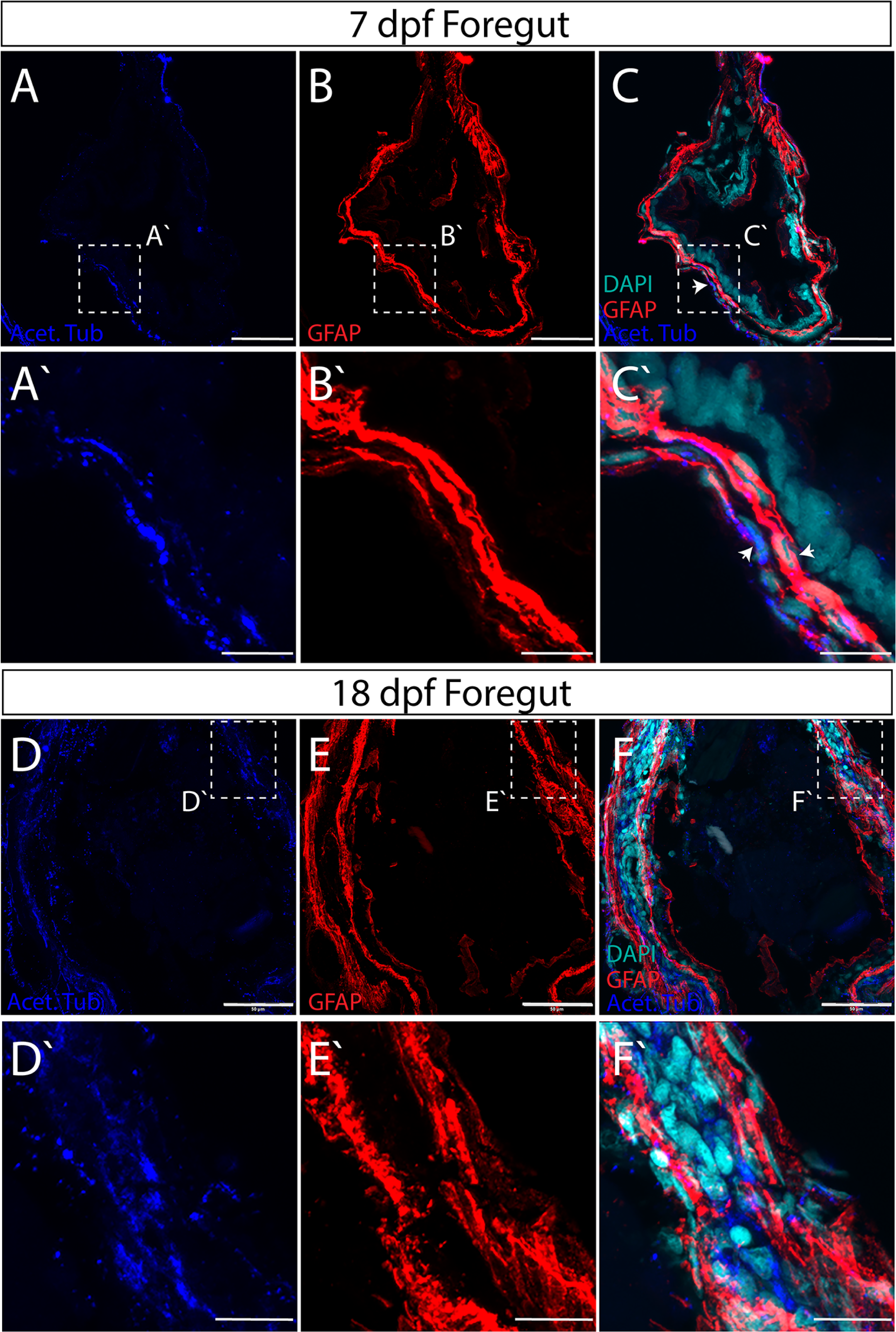
Axon and glial cell patterning within the myenteric plexus of the larval zebrafish foregut. Maximum intensity confocal projections of transverse cryosections indicate GFAP^+^ (red), Acetylated Tubulin^+^ (Acet-Tub) projections (blue) in the foregut of 7 dpf larvae (A-C), where Acet-Tub^+^ processes form in an outer layer (white arrowheads), scale bar: 40 microns, (D-F) 18 dpf larvae, scale bar: 50 microns. Nuclei revealed by DAPI (cyan). White-dashed box corresponds to region of magnification (A`-C`) scale bar: 10 microns, and (D`-F`) scale bar: 12.5 microns.

An increase in overall gut size and complexity occurs between 7 and 18 dpf larval development, which was also exemplified in the ENS neuropil. Within the foregut and hindgut in particular, a notable neuropil layer expansion occurs between 7 and 18 dpf (Fig.3; Fig.5). In both regions, glial and axonal layers expand in width and complexity. For example, within the foregut an additional distinct layer of GFAP^+^ glial processes form on the outer rim of the myenteric plexus by 18 dpf, effectively sandwiching a layer of Acet-Tub^+^ axons (Fig.3E-F’). Furthermore, GFAP^+^ projections interweave throughout the inner plexus neuropil layer of the midgut and hindgut (Fig.4E; Fig.5E).

**Figure 4:**
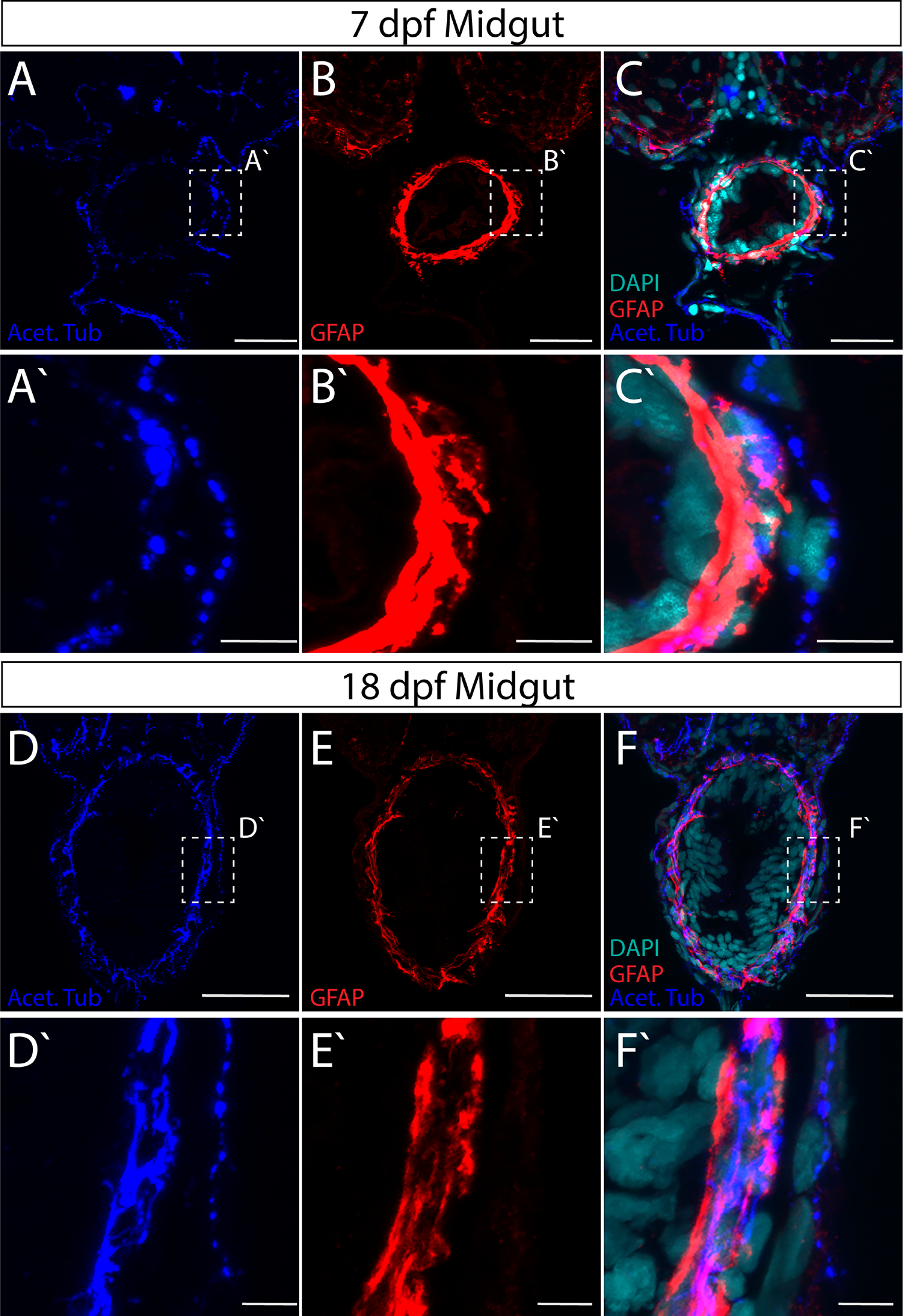
Axon and glial cell patterning within the myenteric plexus of the larval zebrafish midgut. Maximum intensity confocal projections of transverse cryosections indicate GFAP^+^ (red) and Acet-Tub^+^ projections (blue) in in the midgut of (A-C) 7 dpf larvae, scale bar: 20 microns, and (D-F) 18 dpf larvae, scale bar: 40 microns. Nuclei revealed by DAPI (cyan). White-dashed box corresponds to region of magnification (A`-C`) scale bar: 5 microns, and (D`-F`) scale bar: 5 microns.

**Figure 5:**
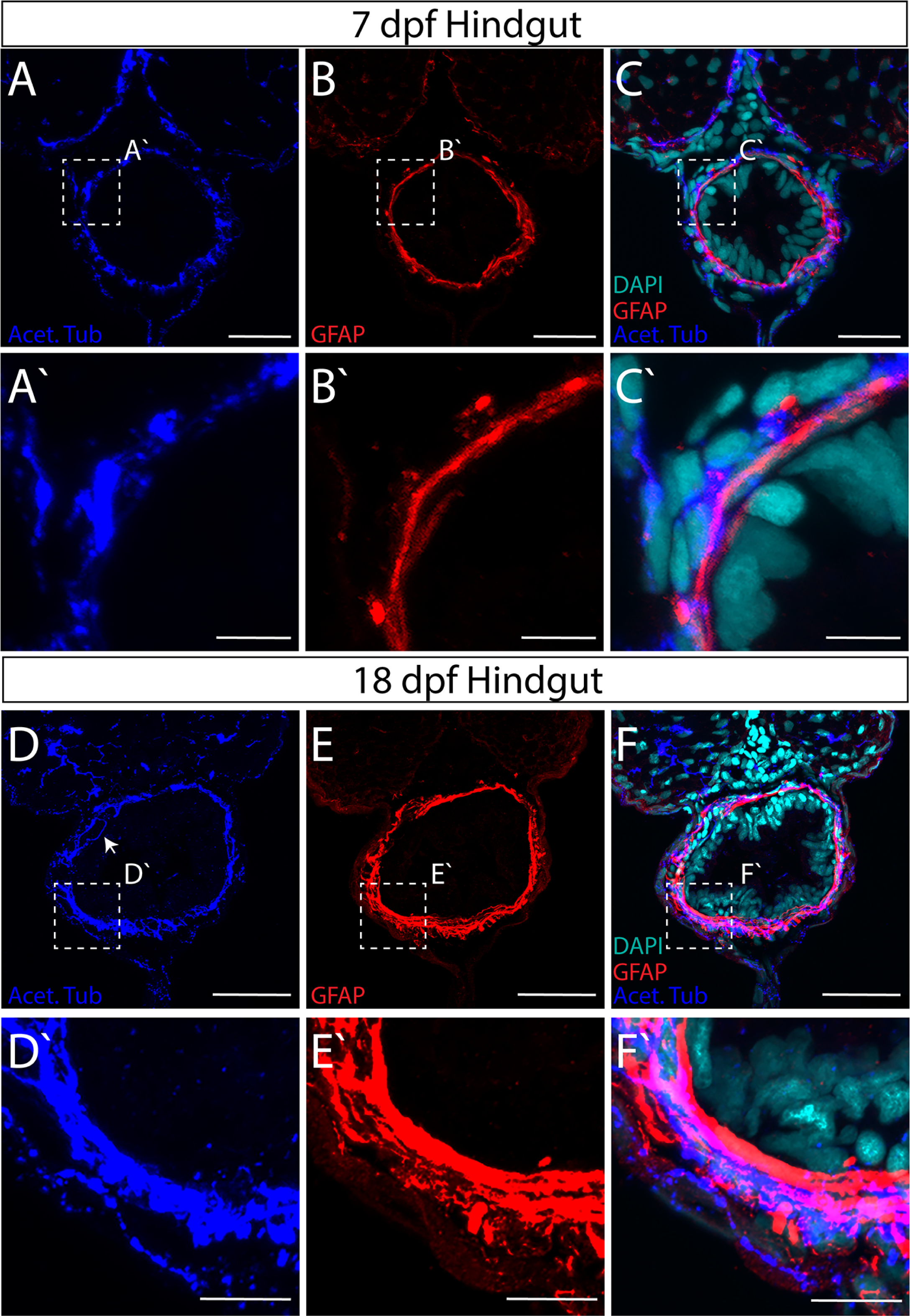
Axon and glial cell patterning within the myenteric plexus of the larval zebrafish hindgut. Maximum intensity confocal projections of transverse cryosections indicate GFAP^+^ (red) and Acet-Tub^+^ projections (blue) in the hindgut of (A-C) 7 dpf larvae, scale bar: 20 microns, and (D-F) 18 dpf larvae with axon projecting close to IE (arrow), scale bar: 40 microns. Nuclei revealed by DAPI (cyan). White-dashed box corresponds to region of magnification (A`-C`) scale bar: 5 microns, and (D`-F`) scale bar: 5 microns.

Based on this analysis, we find that throughout larval maturation, the myenteric plexus is organized in concentric layers of glial cell and axonal neuropil layers of various complexities. These data indicate that the maturing ENS exhibits developmental growth and plexus patterning between 7 and 18 dpf, suggesting that fine cellular interactions underlie the final construction of the mature zebrafish ENS.

### III. Ultrastructural Organization of the myenteric plexus in the maturing larval zebrafish

To investigate the myenteric plexuses we identified in our immunohistochemistry on an ultrastructural level, we utilized transmission electron microscopy (TEM). TEM analysis reveals a neuropil layer, where glial cells and axons are apparent, at 7 and 18 dpf along all levels of the gut (Fig.6-8). In the foregut myenteric plexus at 7 dpf, just below the IE and smooth muscle (M) layer, varicose axons are observed containing many large granular vesicles (Fig.6A, arrows), as has similarly been seen in myenteric plexus of the teleost fish species *Myoxocephalus*^26^ and in adult zebrafish^27^. Immediately juxtaposed to axon profiles are large glial cell bodies (Fig.6A,A’), identified by ultrastructural characteristics of their cytoplasm in cell body and cellular processes. Previously, glial cells have been described based upon the presence of glial filaments (“gliofilaments”)^16^ that exhibit either a filamentous appearance, Type 1 (T1G), or glial cells that exhibit a less filamentous appearance, Type 2 (T2G)^18^. In the larval ENS, we identify cells that resemble both types along all levels of the muscularis (Fig.6-8). For T1G, the glial filaments may exhibit a granular appearance (Fig.8B’,E) or a filamentous appearance (Fig.6B’; Fig.7B’). For T2G, their cellular processes make contacts with T1G and may exhibit areas of local membrane densities (for ex. Fig.7A’,B’, arrows).

**Figure 6:**
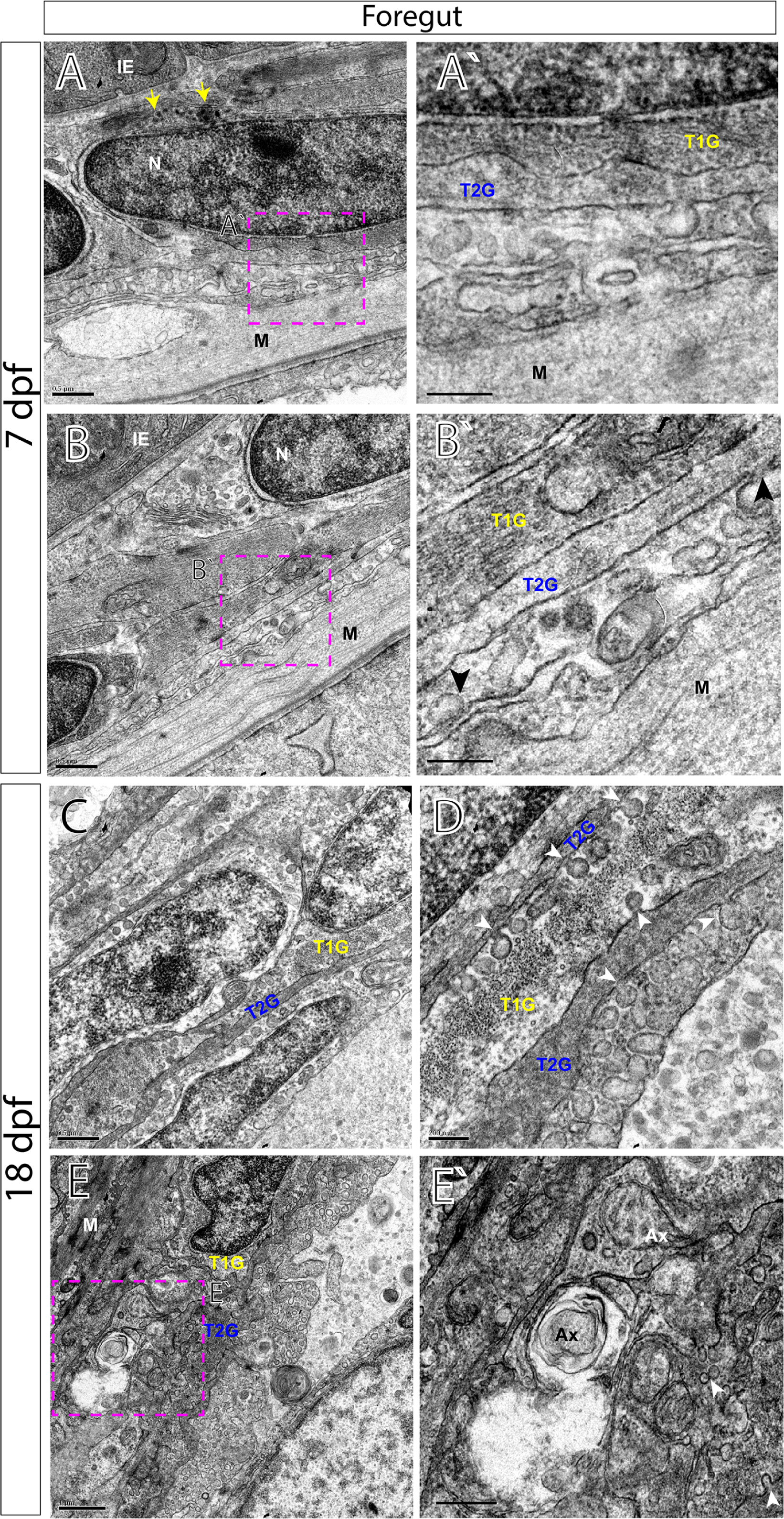
Transmission electron microscopy (TEM) characterizes glial cells and axon ultrastructure within the larval zebrafish foregut. TEM reveals ultrastructure of myenteric plexus neuropil of the foregut in (A-B`) 7 dpf and (C-E`) 18 dpf larvae. Magenta-dashed box corresponds to region of magnification (A`-E`). Intestinal epithelium (IE), nucleus (N), nuclear body (NB), muscularis (M), axon (Ax), type 1 glia (T1G) and type 2 glia (T2G). Scale bars denote the following: (A) 500 nm, (A`) 250 nm, (B) 500 nm, (B`) 250 nm, (C) 500 nm, (D) 200 nm, (E) 1 micron and (E`) 500 nm. Yellow arrows in (A) point to large granular vesicles, black arrowheads in (B’) and white arrowheads in (D,E`) point to caveolae.

**Figure 7:**
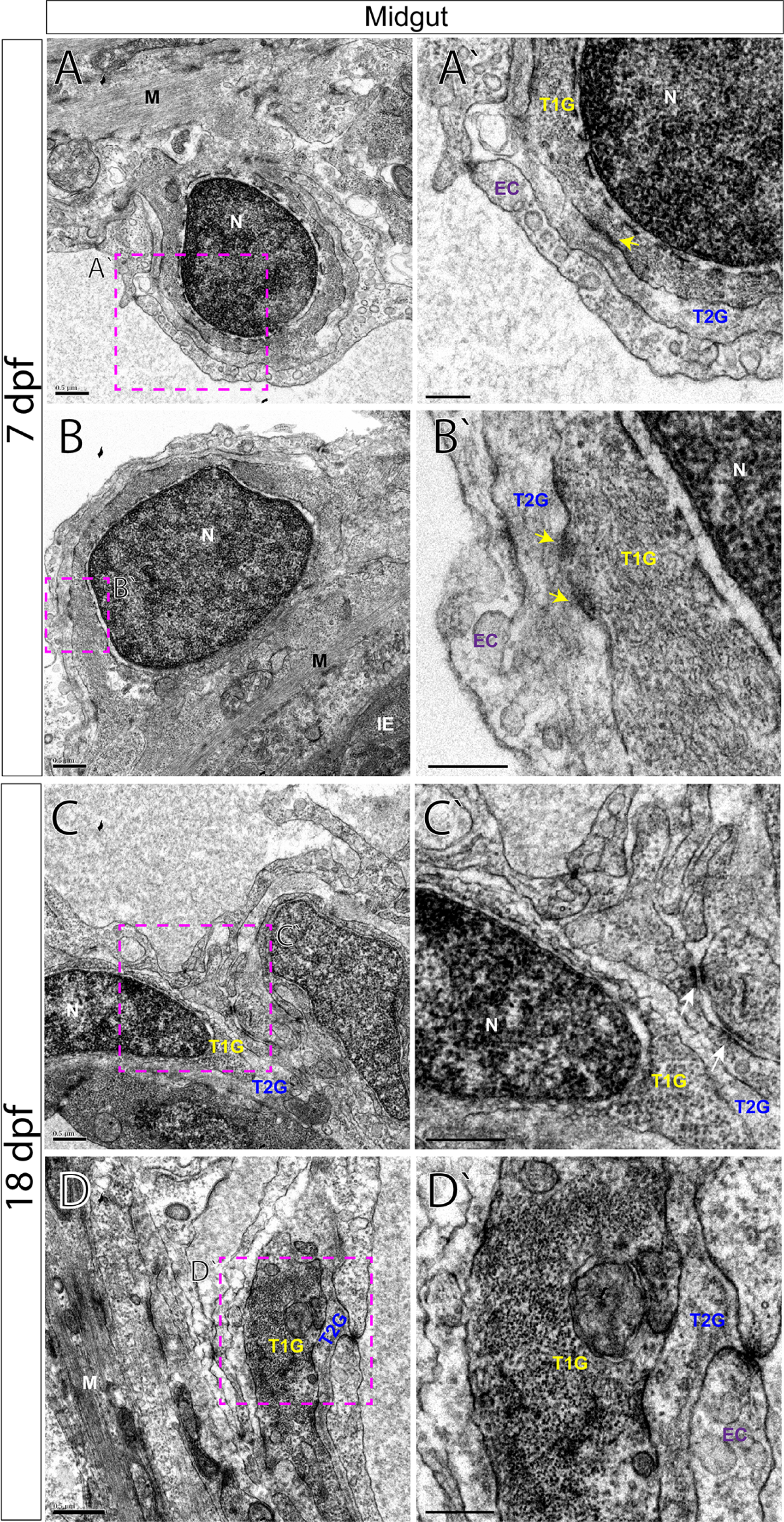
TEM characterizes glial cells and axon ultrastructure within the larval zebrafish midgut. TEM reveals ultrastructure of myenteric plexus neuropil of the midgut in (A-B`) 7 dpf and (C-D`) 18 dpf larvae. Magenta-dashed box corresponds to region of magnification (A`-D`). Intestinal epithelium (IE), nucleus (N), muscularis (M), endothelial cell (EC), type 1 glial (T1G) and type 2 glia (T2G). Scale bars denote the following: (A) 500 nm, (A`) 250 nm, (B) 500 nm, (B`) 250 nm, (C) 500 nm, (C`) 250 nm, (D) 500 nm, (D`) 250 nm. Yellow arrows in (A`,B`) and white arrows in (C`) point to electron dense junctions between T1G and T2G.

**Figure 8:**
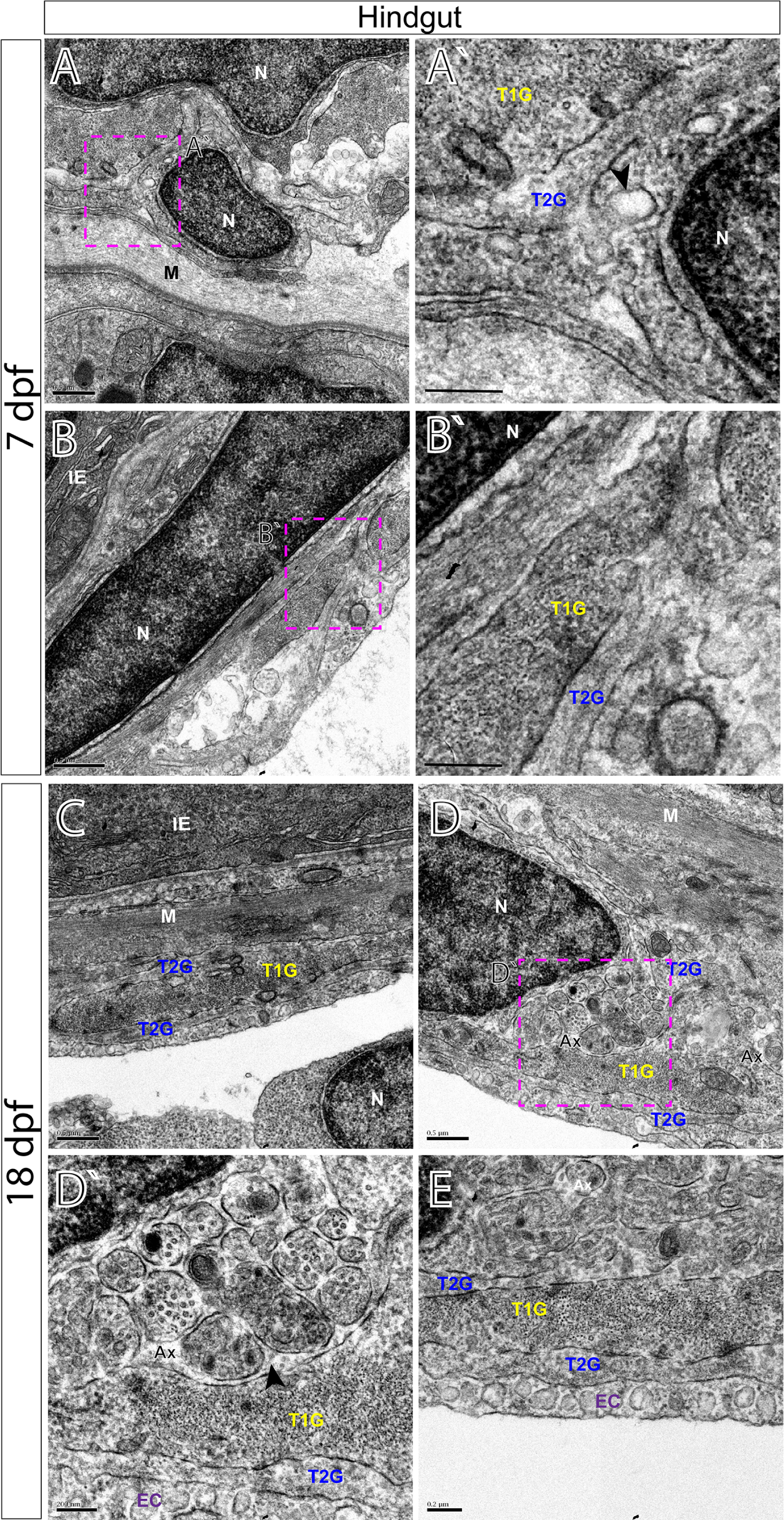
TEM characterizes glial cells and ultrastructure within the larval zebrafish hindgut. TEM reveals ultrastructure of myenteric plexus neuropil of the hindgut in (A-B`) 7 dpf and (C-E) 18 dpf larvae. Magenta-dashed box corresponds to region of magnification (A`-D`). Intestinal epithelium (IE), nucleus (N), muscularis (M), axon (Ax), endothelial cell (EC), type 1 glial (T1G) and type 2 glia (T2G). Scale bars denote the following: (A) 500 nm, (A`) 250 nm, (B) 500 nm, (B`) 250 nm, (C) 500 nm, (D) 500 nm, (D`) 200 nm and (E) 200 nm. Black arrowhead in (A`) points to caveolae and to axon bundles in (D`).

All glial cells present thin filiform processes that wrap the circumference of the muscularis and make direct contacts with each other in parallel tracks at both 7 and 18 dpf to completely ensheathe the gut, in agreement with our immunohistochemical analysis. For example, along the foregut, T2G run parallel to T1G in tight association along long tracks (Fig.6A-B’). A similar pattern is seen along the midgut (Fig.7B,B’) and hindgut (Fig.8C,E). Additionally, we observe that endocytic vesicles, or caveolae, are sometimes shared between filiform glial processes, often “budding” off and in between each other along the foregut (Fig.6D). These glial vesicles exhibit various electron densities, with some being more electron-dense and some are electron-lucent, for example (Fig.6D, arrowheads). Abundant caveolae are also present along an outer layer cellular process, resembling endothelial cells (EC)^28^ which make direct contact with lateral glial cell processes along all levels of the gut (Fig.6D; Fig.7A’; Fig.8E).

By 18 dpf, axon body profiles (Fig.6C) and numerous axon bundles are observed of various densities surrounded by glial cells, indicating that neuronal organization has matured between 7 and 18 dpf (Fig.6E’; Fig.8D,D’). Neuron bodies (NB) display nuclei with a prominent nucleolus, accompanied by electro-lucent cytoplasm (Fig.6C). In the hindgut myenteric plexus, clusters of axon bundles ranging in number from 4-20 are often noted (Fig.8D,D’). The axons range in size from 100 to 400 nm in diameter. The axon profiles contain vesicles of many densities and sizes. Previously, in the mammalian myenteric plexus, varied nerve profiles have been classified into six major types, based upon their ultrastructural morphologies and densities^17,29^. In essence, these include Small Granular Vesicles (SGV), Small Agranular Vesicles (AGV), Small Flattened Vesicles (SFV), Heterogeneous Granular Vesicles (HGV), Large Opaque Vesicles (LOV) and Large Granular Vesicles (LGV)^17^. In the zebrafish larval enteric plexus, we readily find evidence for SGV, HGV, LOV and LGV vesicles within axon bundle clusters at 18 dpf (Fig.8D’). Overall, clear agranular small vesicles, large semi-opaque and dense-core vesicles are all easily seen. These observations demonstrate that zebrafish contain axon profiles similar to those seen in mammalian ENS.

## Discussion

This report describes histology of the zebrafish larval myenteric plexus in high resolution. These data provide baseline characterization of the maturing enteric neuropil along all levels of the gut, thus allowing for fine comparative analysis of phenotypic and/or mutant states moving forward. Zebrafish may provide a fruitful, simplified vertebrate model of the maturing ENS, during stages analogous to post-natal development in mammals. Indeed, it has recently been appreciated that the ENS undergoes continued growth, patterning and maturation during post-natal stages^30–32^, highlighting the need for robust, genetically tractable model organisms to closely examine these phenomena *in vivo*. When combined with stellar live imaging, genetic manipulation attributes and high clutch numbers, zebrafish make excellent candidates for future ENS research.

Collectively, our immunohistochemical and ultrastructural detection of axonal and glial cell profiles in the larval gut reveals novel details about zebrafish ENS architecture and maturation. We observe that the zebrafish larval enteric plexus is histologically distinct by 7 dpf and contains axon and glial cell profiles that encircle the intestinal epithelium. This plexus contains a neuropil, evident by the presence of an axonal layer immediately juxtaposed to glial cells—along the foregut, midgut and hindgut (Fig.9A,B). Previously, it has been appreciated that the zebrafish ENS contains neurons that are sparsely distributed along the surface of the gut tube during larval stages, as well as in the adult^21,22^. From these previous studies we know that a simple meshwork of enteric neurons interconnect in a web-like structure to coat the gut tube. Zebrafish enteric neurons do not cluster to form ganglia, as is seen in mammals^1,12^. Nonetheless, in zebrafish, the enteric neurons express combinations of neurochemical markers; including serotonin (5HT), tyrosine hydroxylase (TH), pituitary adenylate cyclase-activating peptide (PACAP), vasoactive intestinal peptide (VIP), choline acetyltransferase (ChAT) and nitric oxide (NO)^22^, demonstrating that zebrafish use conserved neurochemicals abundant in mammalian nerve profiles of the ENS^1^. Therefore, in building upon these previous studies, our description of an enteric neuropil extends knowledge of the anatomy of zebrafish plexus architecture.

**Figure 9:**
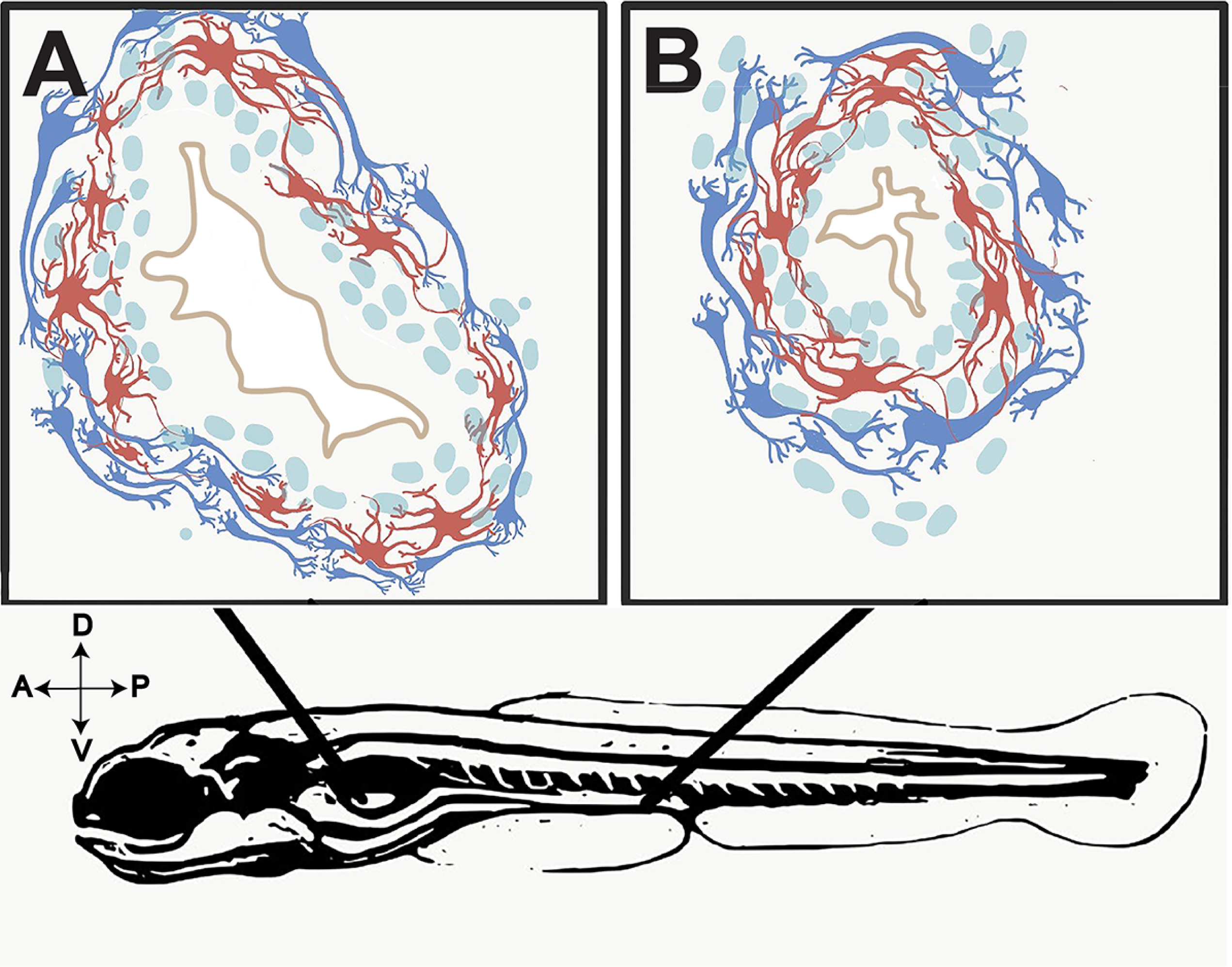
Illustrative representation of ENS myenteric plexus structure in the maturing larval zebrafish. Representative illustrations of (A,B) immunohistochemical preparations of 7 dpf larval foregut and hindgut depicting glia (red) and neurons (blue) of the myenteric plexus. Not to scale.

The complexity of the zebrafish enteric plexus increases by 18 dpf, highlighted by expansion of an outer glial cell layer and thickening of the axon layer. Regionally, we find that the foregut displays the most complex pattern maturation, exhibiting the greatest thickness of myenteric plexus observed (Fig.3). The midgut and hindgut also undergo an expansion in size between 7 and 18 dpf (Fig.4; Fig.5). In the human gut, it is known that the myenteric plexus undergoes extensive growth and patterning between weeks 10 and 18 of gestation, during which time the fetal gut exhibits motor activity^33^. Over time, myenteric plexus nerve profiles in teleost fishes encompass those of the essential neural reflexes of the gut—including motor neuron, interneuron and sensory^1,34^. That we observe the expansion in myenteric plexus complexity between 7 and 18 dpf suggests such fine neural reflex patterning may be actively occurring during this developmental stage. Accordingly, we note in our ultrastructural analysis, the presence of axon bundles distributed longitudinally throughout the enteric plexus by 18 dpf (Fig.8), further supporting the idea that ENS maturation is active during late larval stages.

In corroboration with our immunohistochemical analysis, TEM provides evidence of axon profiles and glial cell bodies distributed throughout the neuropil in 7 and 18 dpf larval zebrafish. Our investigation of the foregut, midgut and hindgut of 7 dpf and 18 dpf fishes reveal a common pattern that is prevalent throughout the larval zebrafish: long tracks of glial cell fibers that lie in close association with one another (T1G, T2G), which cover the extent of the gut tube to completely encapsulate it. While enteric glial cells have been described in the zebrafish gut during early larval stages (3-9dpf) and in the adult fish in whole mount preparations^35–37^, their localization in older larval fish and their histological distribution in the plexus was unknown. In our observations, we note that glial cells are abundant in the maturing larval gut. In fact, we also observe that glial cell processes may exchange matter between one another, in the form of vesicle transfer, suggested by caveolae (Fig.6D). It has been supposed that astrocytes, a type of glial cell, may participate in “gliotransmission”, via release of neuroactive substances in order to modulate neural activity^38^, raising the possibility that a similar mechanism may exist in the gut. Moreover, we find evidence of axon types in zebrafish, similar to those seen in mammals. In the primate and guinea pig myenteric plexus, for example, it is known that several classes of vesicles demarcate particular axonal profiles; notably that agranular vesicles represent cholinergic varicosities, dense-core (osmiophilic) granules suggest adrenergic and that large vesicles with medium electron density denote the “P-type”, peptide-containing vesicle^39^. Based on these descriptions, our data collectively suggests that zebrafish display conserved indicators of these profiles and a maturing myenteric plexus.

In summation, study of the zebrafish larval enteric plexus is timely. Our descriptions here extend knowledge of structure and organization of the zebrafish myenteric plexus, in high resolution. They reveal that during larval maturation stages, the neuropil undergoes a growth phase, highlighted by the manifestation of increased complexity by 18 dpf. In the future, these histological descriptions will provide baseline data with which to compare zebrafish models harboring mutations or gene knockouts in candidate genes of interest that may play a role in ENS construction and/or function.

## Materials and Methods

### Zebrafish Maintenance and lines

Zebrafish (*Danio rerio*) were maintained at 28.5°C on a 13-hour light/11 hour dark cycle. Animals were treated, and experiments performed, in accordance with Rice University IACUC (Institutional Animal Care and Use Committee) approved provisions and guidelines, protocol number 1143754. Wild-type AB (Zebrafish International Resource Center) at 7 and 18 dpf were used for this study.

### Histology and Transmission Electron Microscopy

Freshly euthanized larvae were fixed overnight in Karnovsky’s fixative^40^, rinsed in 0.1M sodium cacodylate buffer, post-fixed for one hour in 1% osmium tetroxide, then again rinsed in 0.1M sodium cacodylate buffer. Larvae were then dehydrated in a graded series of ethanol, infiltrated with and embedded in epoxy resin, and polymerized in a 70 °C oven for 48 hours. To identify areas of interest for TEM analysis, semi-thin sections of 1 µm thickness were cut transversely using an ultramicrotome, stained with toluidine blue and imaged with a Nikon Instruments Eclipse compound light microscope. Ultra-thin sections of 100 nm thickness for TEM analysis were then cut using an ultramicrotome and stained with 33% methanolic uranyl acetate for 15 minutes and lead citrate (Electron Microscopy Sciences, cat # 22410) for 7 minutes. Digital electron micrographs were collected using a JEOL JEM-1230 equipped with a Gatan CCD camera.

### Immunohistochemistry

Immunohistochemical preparations were performed on larvae that were fixed and treated as previously described^7^. All primary and secondary antibody solutions were made using 5% goat serum block in PBTD (1% DMSO, 1X Phosphate Buffered Saline, .1% Tween-20). The primary antibodies were used at the following dilutions: Mouse anti-Acetylated Tubulin 1:200 (Sigma-Aldrich, T7451) and Rabbit anti-GFAP (zebrafish epitope) 1:500 (Genetex, GTX 12874). The secondary antibodies and their dilutions were used as follows: Alexa Fluor 488 Goat anti-Rabbit 1:700 and Alexa Fluor 647 Goat anti-Mouse 1:500 (Invitrogen). DAPI was incorporated into the secondary antibody solutions to mark nuclei following incubation on cryosections.

An Olympus Fluoview FV3000 confocal laser-scanning microscope was used to image the cryosections. Whole-view sections were imaged using an UCPlanFLN 20x/0.70 objective. To capture higher magnification images focused on the gut, an UPlanSApo 60x/1.35 oil immersion objective was used. Z-stacks were obtained by capturing slices every 1-1.5 µm ranging from 10-13 µm total thickness. Images were processed through Olympus CellSen Dimensions software to generate maximum intensity Z-stack projections prior to exporting the images as .tif files. All images were further processed and cropped using Acrobat Adobe Photoshop CC software.

### Data Availability

Data will be made available upon request.

## Acknowledgements

We thank Eileen Singleton for fish care and technical help. We thank Rice University Biosciences department startup funds for funding. R.A.U. is a Cancer Prevention & Research Institute of Texas (CPRIT) Scholar in Cancer Research.

## Author Contributions

R.A.U., M.D.M and P.A.B. designed the study; M.D.M and P.A.B. performed experiments and collected data; R.A.U., P.A.B., M.D.M. and A.T. analyzed data, drafted and edited the manuscript.

